# Bayesian genomic models boost prediction accuracy for resistance against *Streptococcus agalactiae* in Nile tilapia (*Oreochromus nilioticus*)

**DOI:** 10.1101/2020.01.09.900134

**Authors:** Rajesh Joshi, Anders Skaaurd, Alejandro Tola Alvarez, Thomas Moen, Jørgen Ødegård

**Affiliations:** GenoMar Genetics AS, Tjuvholmen allé 11, 0252 Oslo, Norway.; AquaGen AS, P.O. Box 1240, Sluppen, 7462 Trondheim, Norway.

**Keywords:** Genomic selection, Streptococcosis, prediction accuracy, Streptococcus agalactiae, GBLUP, Bayesian, Nile tilapia

## Abstract

Streptococcosis due to Streptococcus agalactiae is a major bacterial disease in Nile tilapia, and development of the resistant genetic strains can be a sustainable approach towards combating this problematic disease. Thus, a controlled disease trial was performed on 120 full-sib families to i) quantify and characterize the potential of genomic selection for S. agalactiae resistance in Nile tilapia and to ii) select the best genomic model and optimal SNP-chip for this trait.

In total, 40 fish per family (15 fish intraperitoneally injected and 25 fish as cohabitants) were selected for the challenge test and mortalities recorded every 3 hours, until no mortalities occurred for a period of 3 consecutive days. Genotypes (50,690 SNPs) and phenotypes (0 for dead and 1 for alive) for 2472 cohabitant fish were available. The pedigree-based analysis utilized a deep pedigree, going 17 generations back in time. Genetic parameters were obtained using various genomic selection models (GBLUP, BayesB, BayesC, BayesR and BayesS) and traditional pedigree-based model (PBLUP). The genomic models were further analyzed using 10 different subsets of SNP-densities for optimum marker density selection. Prediction accuracy and bias were evaluated using 5 replicates of 10-fold cross-validation.

Using an appropriate Bayesian genomic selection model and optimising it for SNP density increased prediction accuracy up to ∼71%, compared to a pedigree-based model. This result is encouraging for practical implementation of genomic selection for S. agalactiae resistance in Nile tilapia breeding programs.

## Introduction

Nile tilapia is an important aquaculture species because of its wide range of trophic and ecological adaptations, which has enabled it to be farmed in different environments across the world. The farming of Nile tilapia is one of the fastest growing aquaculture activities in more than 120 countries, accounting for 5.3% of global aquaculture production in 2017. The species has been ranked 4^th^ among the top ten aquaculture species, in terms of both production quantity and value [1,2]. For the last three decades, the tilapia sector has seen rapid increase (∼11% increase annually) in global production, which is higher than the average growth in other aquaculture species [3,4]. This intensification of tilapia farming in commercial conditions generally results in stress combined with various factors including sub-optimal temperatures, handling, severe crowding, and poor water quality [5]. Furthermore, farmed tilapia are more exposed to various bacterial, viral, fungal and parasitic diseases than free-living tilapia [6].

Streptococcosis, a disease caused by the pathogens *Streptococcus agalactiae* and *Streptococcus iniae*, has been considered one of the most significant bacterial diseases in Nile tilapia based on socio-economic impact and zoonotic potential [7]. The estimated losses were around 250 million USD in 2008 [8]. Among these two Streptococcus species, S. *agalactiae* is most prevalent [9] and it has been shown to cause significant morbidity and mortality [10], with a mortality rate of >50% in acute infections [11]. Symptoms of the infection are lethargy, erratic swimming, hyper-pigmentation of the skin, exophthalmia with haemorrhagic eye, splenomegaly, abdominal distension and diffused haemorrhage in the operculum, around the mouth, anus and base of the fins [8,12,13].

Various short-term strategies to contain *S. agalactiae* using antibiotics and vaccines are used around the world [14–16]. These methods have their own deficiencies, for example: using antibiotics is expensive and complex because of the long withdrawal period and the rising concern of anti-microbial resistance both in fish and humans [6,14,17]. Thus, alternative long-term strategies to control the disease have been sought after [18]. Development of resistant strains of tilapia is one of those sustainable methods, which have been widely and successfully implemented in aquaculture species like Atlantic salmon [19–21], motivating similar development in various species, including *S. agalactiae* in Nile tilapia [22–25].

Classical selection methods for disease resistance, where resistance is assayed in siblings of the breeding candidates, allows utilisation of the between-family variation only. This puts limits on the accuracy of selection, and increases the risk of inbreeding [26]. Using classical selection methods, genetic selection for *S. agalactiae* in GST® Nile tilapia has resulted in strains/products where the risk of mortality is reduced by nearly two thirds compared to the non-selected line [25]. Genomics could increase the rate of genetic improvement further, by allowing utilisation of within-family genetic variation, thereby increasing the accuracy of selection and decreasing the rate of inbreeding. [26,27].

This paper is the part of wider study on the implementation of genomic selection in Nile tilapia breeding programs. The previous studies [28,29] have shown the benefits in using genomic selection for morphological traits in Nile tilapia. And hence, this paper has tried to improve this understanding beyond the morphometric traits, to the disease resistance trait. The objectives of the present study were: i) to quantify and characterize the potential of genomic selection to *S. agalactiae* control in Nile tilapia. ii) to find the SNP-chip marker density which maximises the prediction accuracy for *S. agalactiae* resistance in Nile tilapia and iii) to select the best genomic prediction model for implementation of genomic selection for *S. agalactiae* resistance in Nile tilapia.

## Methods

### 1. Study population

The breeding program for GenoMar Supreme Tilapia (GST®) in the Philippines is a continuation of the Genetically Improved Farmed Tilapia (GIFT) program. The genetic base of GIFT was formed by the systematic admixture of 8 different wild and commercial strains of Nile tilapia [30]. GenoMar bought generation 10 of the GIFT strain and has since been breeding Nile tilapia for growth, fillet yield and robustness [28]. All the samples used in this study originate from 4 out of 8 batches of Generation 27 of the GST® strain.

Each generation of GST® consists of 250 families distributed across 8 batches that follow a revolving breeding scheme where males from batch n are mated to females from batch n-1. In this way, ∼30 families are produced in each batch, significantly reducing the within-batch age difference compared to how it would have been if all 250 families were made as a single batch. The families within each batch were created by mating the selected parents in a 1:1 mating design, where one male and one female were placed in a small breeding hapa. After mating, eggs were collected, and the families were kept separate until hatching.

### 2. Challenge test and Phenotypes

Controlled disease challenge test was performed using *Streptococcus* agalactiae *Ib* – Costa Rica (AL 20109-4) strain. Overall 108 full-sib families from Generation 27 of the GST® strain were challenged in 4 batches. A pre-challenge test was done to determine the LD_50_, the dose that caused 50% mortality, in IP challenged fish (i.e. injection of the pathogen directly in the intra-peritoneal region of the fish). Each family were kept in separate holding tanks to an average weight of 8-10 g, at which time they were moved to the challenge tanks. Forty fish per family was selected and tagged for the challenge test and were placed into each tank as follows: 15 fish as IP injected (0.05 ml of bacterial strain) and 25 fish as cohabitant fish. Mortalities were recorded every 3 hours. The id and a fin clip were immediately collected from each dead fish. The experiment continued until no mortalities occurred for a period of 3 consecutive days. After the experiment, all the fin clips of the surviving cohabitant fish were collected, and the surviving fish were destroyed. The survival phenotype at the end of the experiment was coded as a binary trait; 0 for the dead fish and 1 for the surviving.

### 3. Genotypes

To reduce genotyping cost only the cohabitant fish were genotyped, as they were considered to best mimic the conditions in a disease outbreak in farm conditions. Genomic DNA was isolated from fin clip samples of the 2688 challenge tested cohabitant fish and genotyping was performed using the Onil50 Affymetrix Axiom Custom Array [31]. Thus, the raw dataset contained 58,466 SNPs and 2688 individuals. These genotypes were subjected to different quality control (QC) criteria. Only high quality SNPs, i.e. SNPs identified as PolyHighResolution and NoMinorHomozygous by Affymetrix’s Axiom Analysis Suite software [32], were selected (50,690 loci). Also individuals were removed if their genotyping rate was less than 90%, leaving 2472 samples for the analysis. The average call rate in the remaining samples was 99%.

### 4. Statistical analysis

Pedigree- and genomic based statistical models were used to estimate variance components and heritabilities for *Streptococcus* resistance in Nile tilapia.

#### Pedigree BLUP (PBLUP)

DMU [33] was used to fit a mixed linear univariate PBLUP model, using REML to estimate the variance components and breeding values, described as:

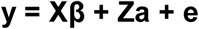

where, **y** is the vector of phenotypes coded as 0 for dead fish and 1 for surviving fish after the challenge test, **β** is the vector of fixed effects that account for batch (3 d.f.), **a** is a vector of random additive genetic effects, **e** is a vector of random residual errors, and **X** and **Z**, are corresponding design matrices for the fixed and random effects. Vectors **a** and **e** had effects for everyone having phenotypes and their distributional assumptions were multivariate normal, with mean zero and

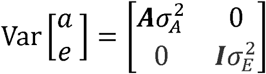

where σ^2^_A_ and σ^2^_E_ are additive genetic variance and error variance respectively; **A** is the numerator relationship matrix and **I** is an identity matrix of appropriate size. The numerator relationship matrix was calculated from pedigree 17 generations back in time. The phenotypic variance was calculated as σ^2^_P_= σ^2^_A_+ σ^2^_E_, and the heritability (h^2^) was calculated as the ratio of the variances (σ^2^_A_/ σ^2^_P_).

#### Genomic Models

Several different types of genomic models are commonly used in animal breeding, including the genomic BLUP (GBLUP) model (either as a marker effects model or as a genomic animal model) and various Bayesian marker effects models (see [34] for detailed explanation). These methods differ primarily in their prior assumptions about the effects of the SNPs.

#### Genomic BLUP (GBLUP)

GBLUP is the most commonly used genomic model for routine genetic evaluations because of its simplicity and less computationally demanding nature. The model can be fitted either as genomic animal model, with SNP-based relationships, or as a marker-effect model, assuming a common normal prior distribution for all markers. These two types of GBLUP models have been proven to be statistically equivalent [35–38].

DMU [33] was used to fit a mixed linear univariate genomic animal model, using REML to estimate the variance components and breeding values, as in pedigree-based (PBLUP) models. The difference between PBLUP and GBLUP is that the numerator relationship matrix in PBLUP model (**A**) is replaced by **G**, the genomic relationship matrix in GBLUP model. The **G** matrix was constructed using VanRaden I [39] as follows:

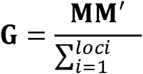

where **M** is a centered marker matrix, the sum in the denominator is over all loci and *p*_*i*_ is the allelic frequency at locus *i*.

The GBLUP model used in this study (using VanRaden I) assumes that all SNP effects to have a common distribution, but the relative contribution of each SNP to the total genetic variance depends on its’ minor allele frequency.

#### Bayesian models

The Bayesian genomic models are less commonly used in routine genetic evaluations due to their more time consuming and computationally demanding requirements. Despite this, it has been shown that they can potentially pick up and use SNPs with large effects or the actual causative variants, especially with disease resistance traits [40,41]. Hence, GCTB2.0 [42] was used to fit four different genomic Bayesian mixed models: BayesB [43], BayesC [44], BayesR [45] and BayesS [42]. All these models were marker effects models, with the general characteristics;

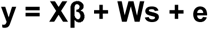

where, **s** is the additive marker locus effect and **W** is the (centred) marker matrix. All other parameters have been described above. All the Bayesian models used in this study are variable selection models, assuming that many of the markers have zero effect and the genetic variation is then explained by a smaller subset of markers. The prior distribution of the variances of **s** differ among these models, as given below:

##### BayesB

Each SNP effect is assumed to have an independent and identically distributed mixture prior of a scaled t-distribution t(0,τ^2^,ν) with a probability π and a point mass at zero with a probability 1-π, where τ^2^ and ν are prior hyperparameters [43,44].

##### BayesC

Each SNP effect is assumed to have an independent and identically distributed mixture prior of a normal distribution having mean 0 and variance σ^2^ with a probability π and a point mass at zero with a probability 1-π [44].

##### BayesR

Each SNP effect is assumed to have an independent and identically distributed mixture prior of multiple normal distributions having mean 0 and variance γ_k_σ^2^_k_ with a probability π_k_ and a point mass at zero with a probability 1-∑_k_π_k_, where γ_k_ is a given constant [45].

##### BayesS

Similar to C but the variance of SNP effects (for loci of non-zero effect) is related to minor allele frequency (p) through a parameter S, i.e. σ^2^_j_=[2p_j_ (1-p_j_)]^S^σ^2^ [42].

All model parameters and SNP effects with the Bayesian models were estimated using the MCMC sampling algorithm implemented in the GCTB2.01 software [42]. The default parameters were used to select the total number of iterations in MCMC (21,000), number of cycles for burn-in (the initial 1000 cycles were discarded) and thinning interval (10). The value of π was estimated from the data using the default starting value of 0.05 (--pi 0.05). Similarly, a default starting value of 0.5 for the sampling of SNP-based heritability (--hsq 0.5) was used.

Like PBLUP and GBLUP, heritability (h^2^) was calculated as the ratio of the additive to phenotypic variances (σ^2^_A_/ σ^2^_P_). PLINKv1.90b6.7 [46] was used to convert the SNP effects obtained from GCTB2.0 software to the breeding values (--score) by simple linear scoring system (i.e. the sum of all the SNP effects).

### 5. Cross-validation and prediction accuracy

The predictive ability of all the models was estimated by using 5 replicates of a 10- fold cross validation scheme. In a 10-fold cross validation, the phenotypes of 10% of animals are masked, and then estimated using the phenotypes and genotypes of the remaining 90% animals. The dataset was randomly divided into 10 sub-sets, predicting one sub-set at a time using the phenotypes of the remaining 9 sub-sets.

Predictive ability of the models was calculated as the Pearson’s correlation between (G)EBVs of all predicted phenotypes adjusted for the fixed effects in one replicate. Results were averaged over the 5 replicates. The obtained mean value of correlation was converted to the expected prediction accuracy by dividing the correlation coefficient by the square root of the estimated heritability (0.15). Standard error of prediction accuracy was calculated [47] as;

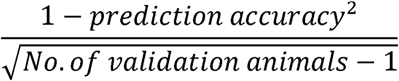

In addition, regression coefficient of fixed effects corrected phenotypes on GEBVs were used to assess the bias of the prediction. The mean value and standard error of the mean regression coefficient were calculated from the five replicates. A regression coefficient of 1 indicates unbiased prediction, whereas the value <1 indicates inflation of (G)EBV and >1 indicates deflation of (G)EBV.

### 6. Low-density SNP subsets

A total of 10 different sub-sets of the SNP panel were also created to assess the potential for using a less dense SNP-set. Generally, SNP selection for a low density SNP chip should aim for at least one SNP to be in linkage disequilibrium (LD) with each QTL of the corresponding trait. Thus, LD based SNP pruning method was used to select different sub-set of SNPs.

The set with all 50,690 SNPs was termed as “All SNPs” panel. In “only LG” sub-set, only the SNPs assigned to specific linkage groups [31] were used. That is, the SNPs not assigned to any linkage groups, and those assigned to the mitochondrial genome [48], were removed. Further, these SNPs on “only LG” subset were pruned based on different LD values in PLINKv1.90b6.7 [49]. The LD values used were 0.1, 0.2, 0.3, 0.4, 0.5, 0.6, 0.7, 0.8 and 0.9 and the subsets are named based on this pruning values. For example: in subset “LD0.1” all SNPs having LD values greater than 0.1 were trimmed. The number of SNPs available for analysis in all the sub-sets are shown in Table 1.

**Table 1:** Total number of SNPs in each sub-set after quality filtering. “All SNPs” includes the SNPs not assigned to the linkage groups, whereas, “Only LG” represent the sub-set with the SNPs in linkage groups only. The columns LD0.1 to LD0.9 represent the subsets obtained after pruning the SNPs based on LD values.

| **LG** | **LD0.1** | **LD0.2** | **LD0.3** | **LD0.4** | **LD0.5** | **LD0.6** | **LD0.7** | **LD0.8** | **LD0.9** | **Only LG** | **All SNPs** |
| --- | --- | --- | --- | --- | --- | --- | --- | --- | --- | --- | --- |
| **1** | 21 | 55 | 146 | 254 | 429 | 649 | 931 | 1234 | 1585 | 2553 | 2631 |
| **2** | 21 | 72 | 160 | 342 | 539 | 769 | 1000 | 1225 | 1497 | 2117 | 2187 |
| **3** | 34 | 94 | 203 | 341 | 538 | 732 | 966 | 1226 | 1432 | 1689 | 1730 |
| **4** | 28 | 78 | 166 | 270 | 406 | 598 | 818 | 1053 | 1337 | 2043 | 2091 |
| **5** | 24 | 62 | 147 | 302 | 488 | 728 | 984 | 1322 | 1659 | 2333 | 2460 |
| **6** | 22 | 46 | 82 | 135 | 237 | 380 | 574 | 812 | 1205 | 2431 | 2557 |
| **7** | 36 | 88 | 191 | 360 | 591 | 898 | 1326 | 1833 | 2466 | 4150 | 4306 |
| **8** | 30 | 88 | 178 | 330 | 502 | 700 | 882 | 1048 | 1212 | 1978 | 2034 |
| **9** | 30 | 67 | 143 | 260 | 409 | 583 | 775 | 982 | 1229 | 1834 | 1874 |
| **10** | 28 | 76 | 147 | 259 | 378 | 508 | 664 | 840 | 1034 | 1714 | 1746 |
| **11** | 34 | 94 | 223 | 418 | 616 | 841 | 1085 | 1325 | 1583 | 2282 | 2394 |
| **12** | 27 | 80 | 160 | 288 | 495 | 708 | 1027 | 1325 | 1671 | 2427 | 2487 |
| **13** | 21 | 76 | 178 | 329 | 507 | 700 | 925 | 1147 | 1382 | 2051 | 2124 |
| **14** | 24 | 65 | 147 | 284 | 510 | 779 | 1065 | 1372 | 1642 | 2428 | 2490 |
| **15** | 25 | 72 | 140 | 253 | 388 | 556 | 732 | 966 | 1234 | 2231 | 2283 |
| **16** | 18 | 40 | 92 | 166 | 283 | 454 | 669 | 948 | 1266 | 2201 | 2316 |
| **17** | 31 | 82 | 203 | 360 | 582 | 849 | 1119 | 1404 | 1695 | 2420 | 2455 |
| **18** | 19 | 43 | 98 | 226 | 365 | 534 | 773 | 1024 | 1297 | 1983 | 2058 |
| **19** | 23 | 63 | 152 | 267 | 459 | 667 | 892 | 1154 | 1372 | 1867 | 1947 |
| **20** | 36 | 78 | 156 | 274 | 470 | 702 | 943 | 1224 | 1548 | 2297 | 2364 |
| **22** | 24 | 60 | 149 | 273 | 438 | 656 | 901 | 1163 | 1401 | 1937 | 1989 |
| **23** | 33 | 65 | 123 | 238 | 374 | 572 | 822 | 1066 | 1330 | 1905 | 1956 |
| **Others** |  |  |  |  |  |  |  |  |  |  | 211 |
| **Total** | **589** | **1544** | **3384** | **6229** | **10004** | **14563** | **19873** | **25693** | **32077** | **48871** | **50690** |

The statistical analysis using genomic models, cross validation and prediction accuracy for all the sub-sets were calculated as described in previous sections.

## Results and discussion

To our knowledge, this is the first study on using genomic data to study genetic resistance of Nile tilapia to any disease, quantifying and characterizing the potential of genomic selection to control *S. agalactiae* in Nile tilapia.

The mean mortality % during the challenge test was 60.2%, ranging from 49.5% to 67% in various batches. The Kaplan-Meier curves [50] for the survival function was plotted for the test period to show the cumulative mortality (Figure 1), which clearly shows the differences between the batches.

**Figure 1:**
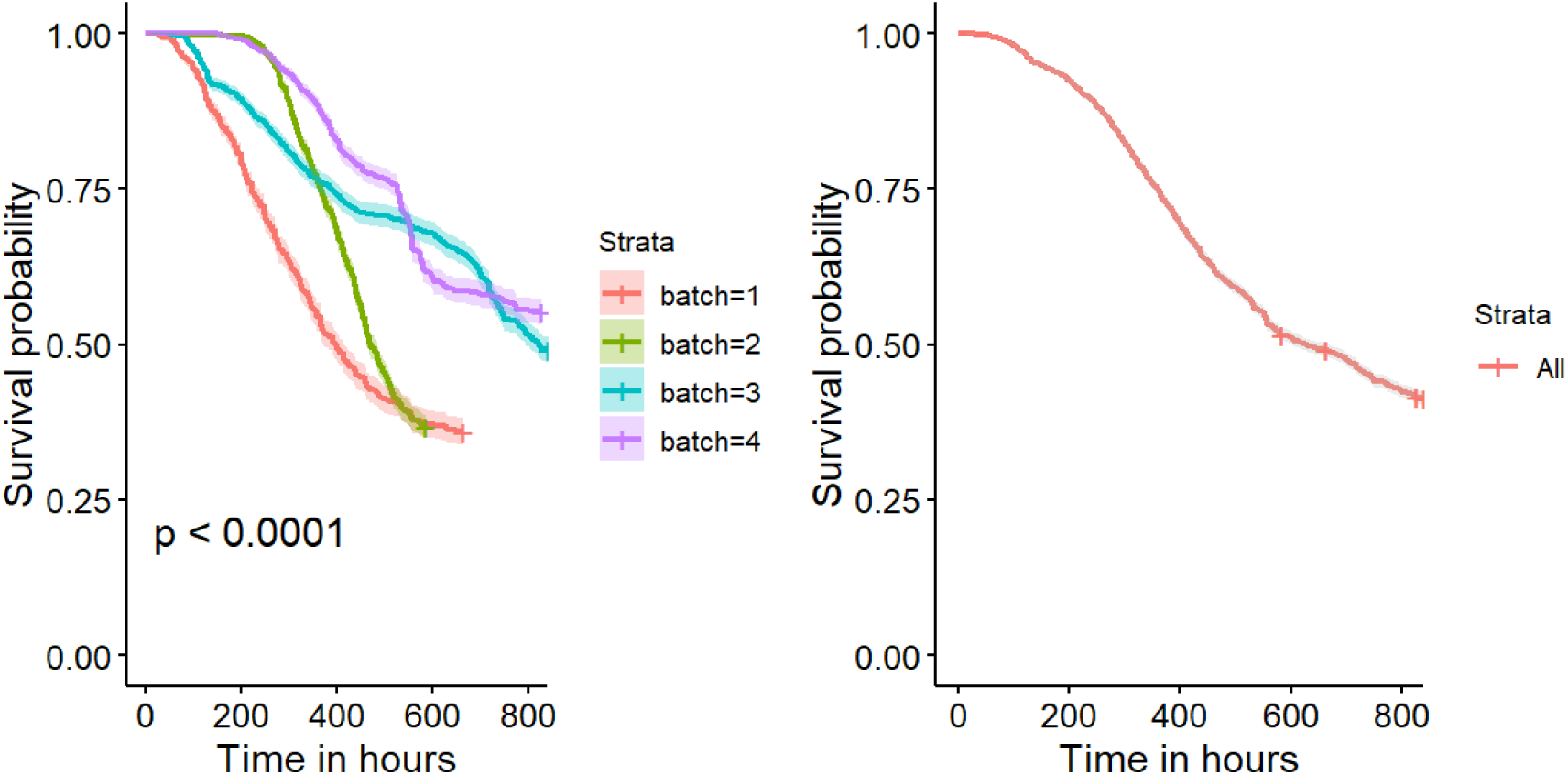
Kaplan-Meier Curves of the survival function for the fish for challenge test. The first figure shows the significant differences between the batches, whereas the second one represents the curve for whole population.

### 1. Genetic parameters

The heritability for *Streptococcus* resistance in Nile tilapia was estimated to be 0.16 ± 0.02 using PBLUP, which is similar to that reported by Sukhavachana et al. [23] and slightly lower than the values reported by Shoemaker et al. [22]. The heritabilities calculated using different genomic models ranged from 0.15 ± 0.03 to 0.26 ± 0.05, as presented in Table 2. Significant estimates of heritability, estimated using the pedigree as well as genomic information, indicate that the Nile tilapia breeding industry can benefit from the application of selective breeding for *Streptococcus* resistance in Nile tilapia.

**Table 2:** Heritabilities for different models and SNP densities for *Streptococcus* resistance in Nile tilapia.

| Sub-set | No. of SNPs | GBLUP |  | BayesB |  | BayesC |  | BayesR |  | BayesS |  |
| --- | --- | --- | --- | --- | --- | --- | --- | --- | --- | --- | --- |
| | | $h^2$ | se | $h^2$ | se | $h^2$ | Se | $h^2$ | se | $h^2$ | se |
| LD0.1 | 589 | 0.09 | 0.02 | 0.06 | 0.01 | 0.09 | 0.02 | 0.10 | 0.02 | 0.10 | 0.02 |
| LD0.2 | 1,544 | 0.16 | 0.03 | 0.17 | 0.02 | 0.18 | 0.02 | 0.17 | 0.02 | 0.18 | 0.02 |
| LD0.3 | 3,384 | 0.16 | 0.03 | 0.17 | 0.02 | 0.18 | 0.02 | 0.17 | 0.02 | 0.19 | 0.02 |
| LD0.4 | 6,229 | 0.19 | 0.03 | 0.22 | 0.02 | 0.21 | 0.03 | 0.20 | 0.03 | 0.23 | 0.02 |
| LD0.5 | 10,004 | 0.19 | 0.03 | 0.22 | 0.03 | 0.21 | 0.03 | 0.19 | 0.03 | 0.23 | 0.02 |
| LD0.6 | 14,563 | 0.19 | 0.03 | 0.22 | 0.03 | 0.21 | 0.03 | 0.19 | 0.03 | 0.23 | 0.03 |
| LD0.7 | 19,873 | 0.19 | 0.03 | 0.25 | 0.03 | 0.23 | 0.03 | 0.19 | 0.03 | 0.23 | 0.03 |
| LD0.8 | 25,693 | 0.18 | 0.03 | 0.24 | 0.03 | 0.25 | 0.02 | 0.19 | 0.03 | 0.23 | 0.03 |
| LD0.9 | 32,077 | 0.17 | 0.03 | 0.23 | 0.03 | 0.26 | 0.03 | 0.17 | 0.02 | 0.23 | 0.03 |
| Only LG | 48,871 | 0.15 | 0.03 | 0.26 | 0.03 | 0.27 | 0.03 | 0.15 | 0.02 | 0.26 | 0.02 |
| All SNPs | 50,690 | 0.15 | 0.03 | 0.26 | 0.03 | 0.25 | 0.05 | 0.26 | 0.05 | 0.24 | 0.04 |

Some studies have claimed that pedigree-based methods overestimate heritabilities [51–53], with other studies suggest the reverse [54–57], and yet others claim heritabilities are similar [58]. Comparing heritabilities coming from pedigree and genomic methods is thus not straightforward, as the methods and models used to calculate the variance parameters and heritability have a direct effect on these parameters [23,59–62], as can also be seen in Table 2.

In all the genomic models, very low SNP densities resulted in lower heritability, as should be expected, as few SNPs are less likely to capture the majority of genetic variance across the genome. For GBLUP, the estimated heritability had a curvilinear relationship with the number of chosen SNPs, with the highest heritability (0.19) found using the moderately pruned SNP-sets (LD0.4 to LD0.7), compared to the highest densities (0.15 for only LG / all SNPs). Increasing marker density should theoretically enable the model to more efficiently capture the majority of genetic variance. In GBLUP, genetic variance is *a priori* assumed evenly distributed over marker loci (depending on the minor allele frequency), but this assumption is only appropriate given that the markers are evenly distributed over the genome. Hence, uneven distribution of marker loci may give a biased representation of the total genetic variance. By pruning SNPs on LD, the number of loci will likely be more evenly distributed over the genome, reducing this bias. For the Bayesian models, reducing the number of SNPs through pruning had largely a negative effect on estimated heritability, compared with using all SNPs. In the Bayesian models the genetic variance of a DNA segment is no longer a function of number of marker loci, but rather depends on the number of included marker loci (and for some of the models, the variance of their effect). Hence, these models are likely more capable of capturing loci of large effect. Further, inflation bias of GEBVs for these models can be seen in Figure 2(b), so the increase in heritability may be bias rather than real.

**Figure 2:**
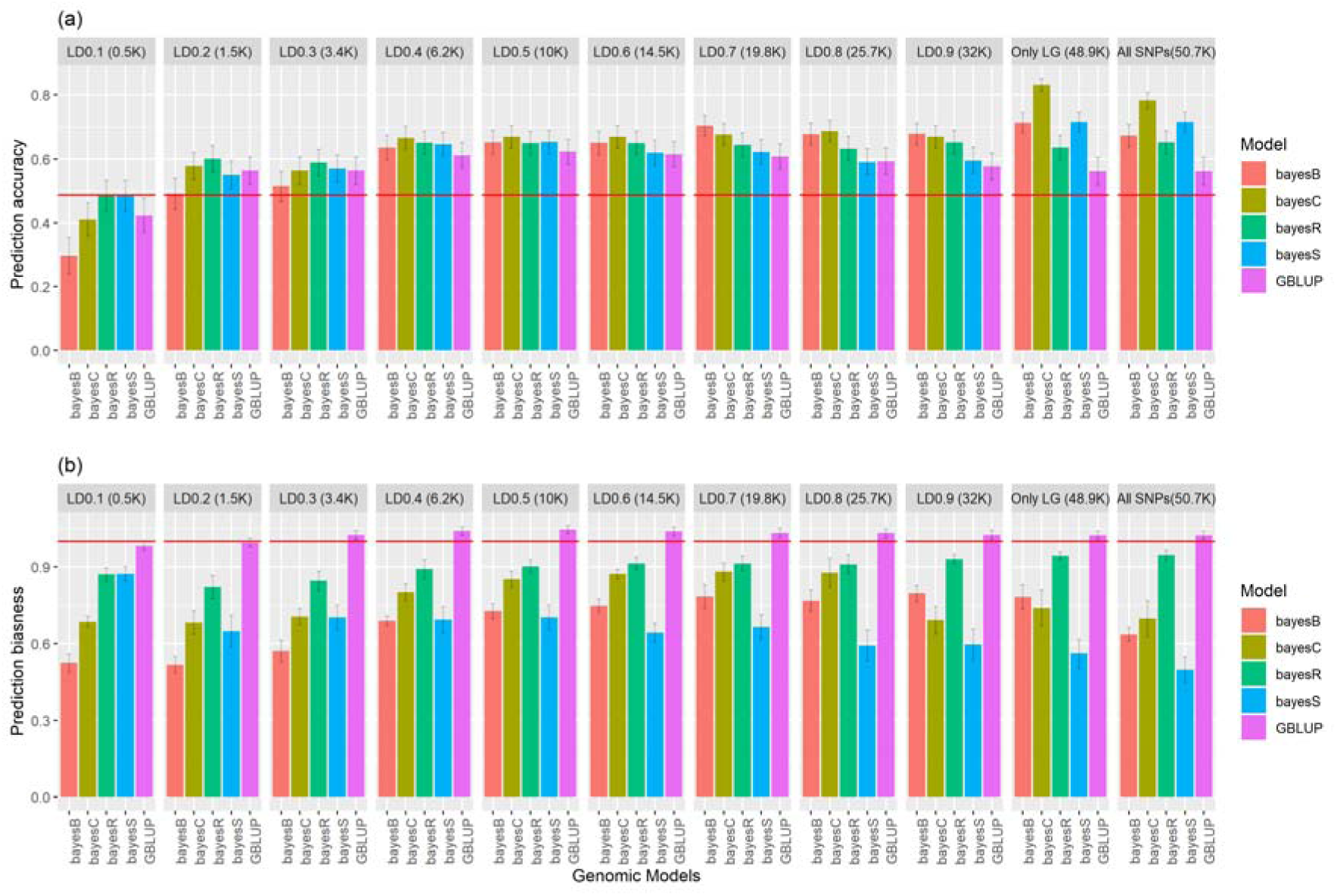
**(a)** Prediction accuracy for *S. agalactiae* resistance in Nile tilapia using different models. The red horizontal line represents the prediction accuracy using PBLUP model. The grey lines in the bar charts represent ± standard errors **(b)** Prediction bias for *S. agalactiae* resistance in Nile tilapia using different models. The red horizontal line represents the prediction biasness using PBLUP model (∼1). The grey lines in the bar charts represent ± standard errors

Another interesting result was to see the impact of excluding the mitochondrial SNPs and SNPs not assigned to any LGs (“All SNPs” vs “Only LG”) on the heritability patterns. The heritability either increased or remained constant using these two SNP sub-sets, but the change was not so marked in almost all the models except BayesR. In BayesR model huge decrease in the heritability estimates was observed using “Only LG” subset of SNP, compared to “All SNPs”.

### 2. Prediction accuracy

The prediction accuracy based on 5-fold random cross-validation, using different models and SNP subsets is shown in Figure 2(a) (see Supplementary Figure S1 for relative increase in prediction accuracy for genomic models, compared to PBLUP). The accuracy of genomic prediction depends on the models applied, which is representative of the architecture of the trait. Thus, using an appropriate genomic model and optimising it for SNP density, we found that the prediction accuracy can be increased by up to ∼71%, compared to a classical pedigree-based model.

#### Optimising genomic models for higher prediction accuracy

The pedigree- and genomic based statistical models used in this study differed in two sources of information: (i) genetic relationship among individuals and (ii) LD between SNPs included. The GBLUP model can exploit realized genomic relationships between sibs, while PBLUP using the expected pedigree relationship [63]. However, compared to GBLUP, Bayesian methods can potentially account for the fact that neither QTL effects nor genotyped SNPs are not necessarily evenly distributed over the genome [64]. Moreover, it should be noted that the training and validation populations used in this study contains closely related individuals, as the relatedness of the training and validation population is directly proportional to the prediction accuracy [65].

##### Advantages of using GBLUP models

As expected, prediction accuracies using the GBLUP model was superior (+15.4%) to the PBLUP model. This increase in prediction accuracy by replacing the pedigree-based numerator relationship matrix by the genomic relationship matrix has been well documented in various aquaculture species (e.g. [41,55,66]). This is due to the fact that GBLUP model can utilise both within- and between-family genetic variation also for traits where the phenotype cannot be measured directly in the selection candidates, like disease resistance [26,27]. The PBLUP model, in contrast, can utilise only between-family genetic variation for such traits.

##### Advantages of using Bayesian models

Prediction accuracy using Bayesian models was found to be superior to both PBLUP and GBLUP models. BayesC was found to give highest prediction accuracy, followed by BayesS, BayesB and BayesR. The advantage of the Bayesian models may be due to the genetic architecture of the studied trait. It has been reported that resistance against *S. iniae* is affected by a major QTL [67] and resistance against *S. agalactiae* is also found to show a similar genetic architecture (results not shown). In GBLUP, all SNPs effects are *a priori* assumed to have the same variance, and their contribution to total genomic relationships thus depends on their MAF only, whereas Bayesian models assumes that the genetic variance is explained by a smaller fraction of the SNPs. Thus, the Bayesian models can potentially pick up and use SNPs of large effect (or the actual causative variants, if included), for example for certain disease resistance traits [40,41], having a few major QTLs (e.g. [68–70]).

##### Low-density SNP panels

The value of constructing low-density SNP panels for routine genomic evaluation has been debated. On one hand, it has been argued that in the species where the economic value per animal is low compared with the cost of genotyping (e.g. Nile tilapia along with other aquaculture species), genotyping large number of selection candidates with high-density SNP panels is not cost effective [71]. Hence, it is desirable to have more cost-effective low-density SNP panels to reduce the cost of genotyping to implement genomic selection in such species. On the other hand, the selection indices for breeding program includes more than one trait (both quantitative and qualitative) and different SNPs are in LD with different QTLs controlling different traits. Hence, multiple low-density SNP panels which are unique for each trait are required, which is not an economical approach. Another approach would be to include all these SNPs in the medium-density SNP panel. But again, the cost of genotyping is decreasing rapidly, and it is not surprising to see very minor differences in genotyping using medium and high-density SNP panels (e.g. 20-30K vs 50K SNP panels). The more realistic cost-effective approach would then be to use low-density SNP panels by imputing it to higher density, but accurate phasing of the genotypes to reduce the error rate require careful planning for high-density genotyping of key ancestors [72].

In general, the SNP data set that maximized estimated heritability for each model also gave the best prediction accuracy (Supplementary Figure S1). Using BayesR and BayesS, we were able to achieve comparable prediction accuracy as PBLUP with only 589 SNPs (smallest subset of SNP panel), whereas a few more SNPs were required for GBLUP and BayesB methods. Hence, these smallest subsets can be cost effective alternatives in cases where pedigree recording is not feasible.

#### Optimising SNP density for higher prediction accuracy

Simulation studies have shown that prediction accuracy decreases gradually with a decrease in SNP panel density [73,74], something which has also been observed in real data [75,76]. We, on the other hand, found that the prediction accuracy increased or remained constant by using only the SNPs in the linkage groups for almost all the models. SNPs not assigned to linkage groups are those SNPs which are not mapped to linkage maps (which may be linked to SNPs genotype quality) and those in mitochondria. After that, the prediction accuracy gradually decreased or again remained constant with decrease in SNP density. These results provide an opportunity to select for minimum SNP density for different models which maximises the prediction accuracy for *S. agalactia* in Nile tilapia breeding programs. The minimum SNP density which could be used for maximising prediction accuracy for GBLUP, BayesB, BayesC, BayesR and BayesS were 10K, 19.8K, 48.9K, 6.2K and 48.9K respectively. Across models, BayesC using 48.9K SNP panel was found to have the best prediction accuracy.

### 3. Prediction bias

The prediction bias using 5-fold random cross-validation, using different models and SNP subsets is shown in Figure 2(b). In Nile tilapia, selection takes place among the single generation of individuals. Hence, selection decision can be made based only in the prediction accuracy, as they share the common mean and bias is not a major concern [77].

Nevertheless, the bias of these prediction accuracies was lowest with GBLUP models in all the datasets. Overall, GBLUP produced largely unbiased predictions, whereas the GEBVs with the Bayesian genomic models were highly inflated (resulting in regression coefficients of predicted phenotyped on GEBVs being < 1). Among the Bayesian models, GEBVs were most inflated for BayesS and least inflated for BayesR for almost all subsets of SNP densities.

## Implications

Nile tilapia aquaculture sector has seen rapid increment in global production for the last three decades and has become one of the most important aquaculture species in the world. This increase in farmed activities have also been followed by increase in disease outbreaks. Breeding for improving genetic resistance can be a way to reduce the huge economic loss due to *S. agalactiae* infection. Thus, using appropriate genomic variable selection model and optimising it for SNP density, we found that the prediction accuracy can be increased by up to ∼71%, when compared to a classical pedigree-based model. Provided that all the managemental practices remain constant, the % of possible increase in genetic gain using genomics is likely even higher, due to less restrictions on inbreeding, compared to classical sib-selection methods. This result is encouraging for practical implementation of genomic selection for *S. agalactiae* resistance in Nile tilapia breeding programs.

## Declarations

### Ethics approval and consent to participate

Not applicable

### Consent for publication

Not applicable

## Availability of data and material

The data used in the study are from commercial family material. This information may be made available to non-competitive interests under conditions specified in a Data Transfer Agreement. Requests to access these datasets should be directed to Alejandro Tola Alvarez.

## Authors’ contributions

RJ conceived the idea, did the statistical analysis and wrote the initial draft of the paper, AA and AS contributed to the design of the project, AS contributed to microsatellite-based pedigree construction, all authors contributed to the discussion of the results and TM and JO contributed to writing of the final version of the paper.

## Acknowledgements

We would like to acknowledge Mayet de Vera for rearing the fish for the experiment.

## Authors’ information (optional)

Not applicable

## Supplementary file

**Figure S1:**
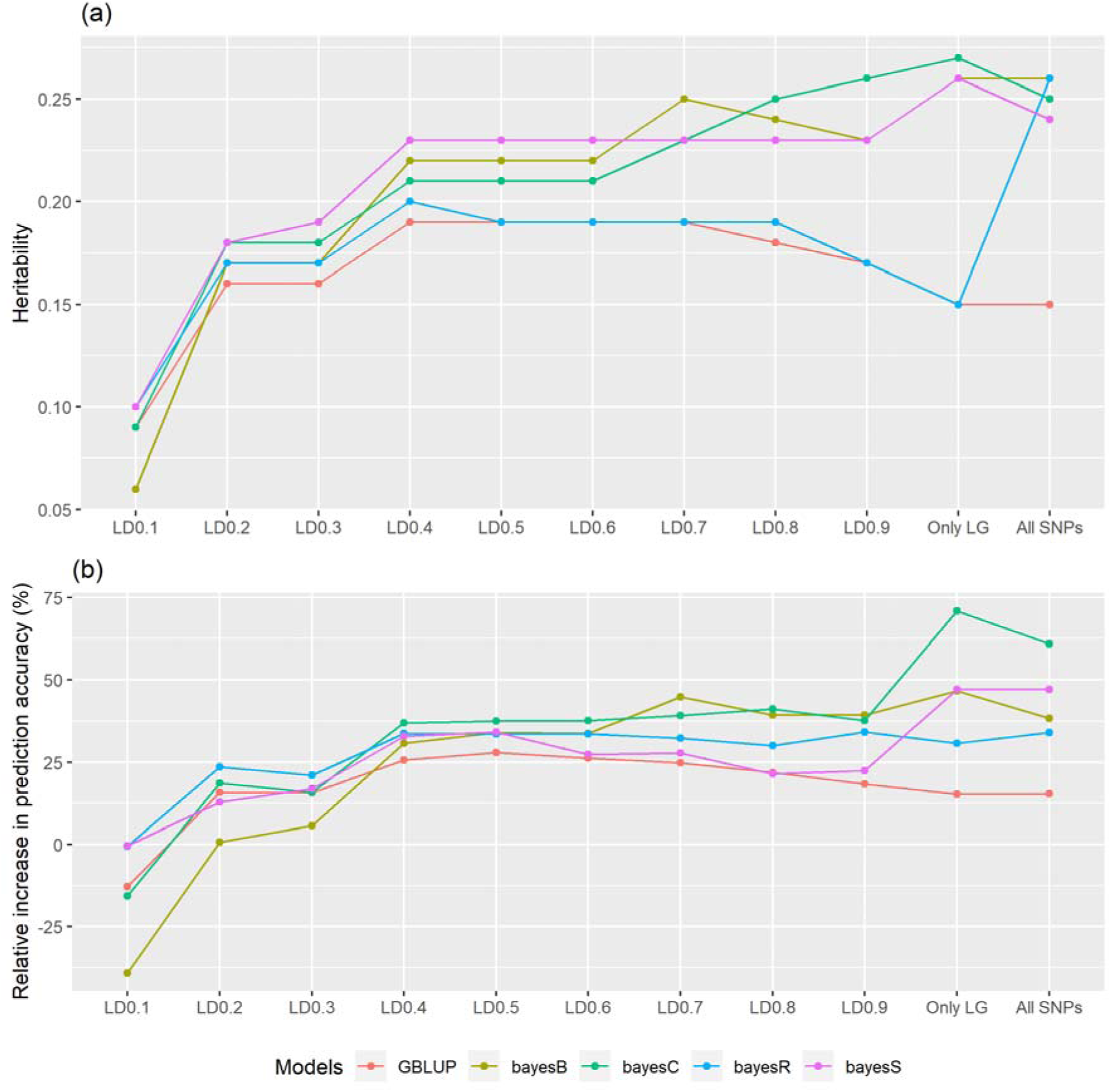
(a) First figure shows the pattern of heritabilities with decreasing SNP density (from right to left) for different models. (b) Second figure shows the pattern of relative increase in prediction accuracy (%) compared to PBLUP with decreasing SNP density (from right to left) for different models.

